# Gene expression landscape of cutaneous squamous cell carcinoma progression

**DOI:** 10.1101/2023.12.11.570862

**Authors:** Tomas Bencomo, Carolyn S. Lee

## Abstract

**Background:** Cutaneous squamous cell carcinomas (SCC) are the second most common human cancer and have been characterized by RNA sequencing (RNA-Seq); however, the transferability of findings from individual studies may be limited by small sample sizes and diverse analysis protocols.

**Objectives:** To define the transcriptome landscape at different stages in the progression of normal skin to SCC through a meta-analysis of publicly available RNA-Seq samples

**Methods:** Whole-transcriptome data from 73 normal skin samples, 46 actinic keratoses (AK), 16 in situ SCC, 13 keratoacanthomas (KA), and 147 SCC (including 30 SCC from immunocompromised patients and 8 SCC from individuals with recessive dystrophic epidermolysis bullosa [RDEB]) was uniformly processed to harmonize gene expression. Differential expression, fusion detection, and cell-type deconvolution analyses were performed.

**Results:** Individual RNA-Seq studies of SCC demonstrated study-specific clustering and varied widely in their differential gene expression detection. Following batch correction, we defined a consensus set of differentially expressed genes (DEGs), including those altered in the preinvasive stages of SCC development, and used single-cell RNA-Seq data to demonstrate that DEGs are often, but not always, expressed by tumor-specific keratinocytes (TSKs). Analysis of the cellular composition of SCC, KA, and RDEB-SCC identified an increase in differentiated keratinocytes in KA, while RDEB-SCC contained the most TSKs. Compared to SCC arising in immunocompetent patients, SCC from immunosuppressed individuals demonstrated fewer memory B cells and CD8 T cells. A comprehensive and unbiased search for fusion transcripts in SCC and intermediate disease stages identified few candidates that recur in >1% of all specimens, suggesting most SCC are not driven by oncogenic gene fusions. Finally, using GTEx data, we distilled a novel 300-gene signature of chronic sun exposure that affirms greater cumulative ultraviolet (UV) exposure in later stages of SCC development.

**Conclusions:** Our results define the gene expression landscape of SCC progression, characterize cell subpopulation heterogeneity in SCC subtypes that contribute to their distinct clinical phenotypes, demonstrate that gene fusions are not a common cause of SCC, and identify UV-responsive genes associated with SCC development.

**What is already known about this topic?:**

- Cutaneous squamous cell carcinoma (SCC) is one of the most common cancers worldwide.
- Several studies have examined gene expression changes in SCC using RNA sequencing (RNA-Seq) but comparison of their results is difficult due to inter-study variation and diverse bioinformatic pipelines and protocols.
- A few gene fusions have been described in SCC, but a comprehensive characterization of fusion transcripts in patient samples has not been performed.

**What does this study add?:**

- We re-analyzed RNA-Seq data from 11 studies of SCC and its preinvasive stages to create a list of consensus differentially expressed genes and identify those that are UV-responsive.
- Clinically aggressive SCC displayed more tumor-specific keratinocytes, while keratoacanthomas contained more differentiated keratinocytes. SCC in immunocompetent persons had more memory B cells and CD8 T cells than those arising in immunosuppressed individuals.
- Previously reported gene fusions were not detected and most fusion candidates did not demonstrate pathogenic features.

**What is the translational message?:**

- Our analysis harmonizes differing results from previous studies to provide a robust list of genes implicated in SCC development.
- Our findings suggest gene fusions are not a common driver event in SCC.

## Introduction

Cutaneous squamous cell carcinoma (SCC) is one of the most common human cancers worldwide (1). Characterization of SCC by microarray-based approaches and more recently by RNA sequencing (RNA-Seq) has provided valuable insight into changing gene expression and an opportunity to detect oncogenic fusion transcripts (2–34). However, individual studies are often hindered by small sample sizes and diverse analysis protocols that can limit the transferability of their results. While meta-analyses can help harmonize differing results between studies (35), a meta-analysis of SCC RNA-Seq has not been performed.

We therefore conducted an analysis of 306 publicly available RNA-Seq samples representing different stages in the progression of normal skin (NS) to SCC. Data from NS, precancerous actinic keratoses (AK), in situ/intraepidermal carcinomas (IEC), keratoacanthomas (KA), and SCC were uniformly processed to remove study-specific batch effects. After harmonization of gene expression, the integrated dataset was used to derive a consensus list of differentially expressed genes and generate a comprehensive catalog of fusion transcripts present in the SCC progression continuum. We also employed this dataset to characterize cell subpopulation heterogeneity in SCC subtypes and highlight, using a novel 300-gene signature of chronic sun exposure, ultraviolet (UV)-responsive genes altered in SCC development.

## Materials and Methods

### Study selection

We performed a literature search for studies with publicly available sample-level RNA-Seq data of SCC tumors and related lesions. Studies were required to have raw FASTQ files available on public databases (SRA, GEO, ArrayExpression, ENA) without restriction and at least 5 million reads per sample.

### Sample Processing

Raw FASTQ files from each study were downloaded from SRA or ArrayExpress and aligned to GENCODE v38 using STAR (v2.7.9). Read counts were quantified with RSEM (v1.3.3). The STAR-Fusion (v1.12.0) pipeline was used for fusion detection. Reads were aggregated into a count matrix with tximport (v1.22.0)(36) and read depth normalized with DESeq2 (v1.34.0).

Count data was transformed using the vst() function in DESeq2 for PCA and UMAP plots as well as other downstream visualization and analyses. For PCA, the top 500 most variable genes were selected for dimensionality reduction. For UMAP, the top 1000 most variable genes were used to compute 50 principal components that were supplied as input to UMAP.

DvP scores were computed by multiplying the DvP signature gene weights with the matching gene’s VST-normalized expression values for each sample, summing the products, and then performing Z-score normalization as previously described (32).

### Differential Expression Analysis

For the individual study analysis, the count matrix for each study was normalized by DESeq2 and the NS vs SCC contrast was computed without adjustment for any factors. P-values were adjusted using the Independent Hypothesis Weighting (IHW, v1.22.0) approach (37) to maximize power and avoid false positives from minimally expressed genes. Genes that were called a DEG with a 5% FDR in ≥7 studies were used as input to EnrichR (v3.2.0) with upregulated and downregulated genes analyzed separately and using the MSigDB Hallmark 2020 genesets. A 10% FDR threshold was used to determine significant pathways.

For the pooled analysis, DESeq2 was run on the entire count matrix with the design formula ∼ study + sample_type to adjust for study membership. P-values were adjusted using the IHW approach described above. A 0.1% FDR threshold was used for the NS vs SCC comparison and a 1% FDR threshold was used to compare NS to AK and AK to SCC due to the smaller sample sizes. GSEA was performed with the fgsea (v1.26.0) (38) R package using Hallmark and Reactome gene sets downloaded from the MSigDB website (39).

### scRNA-Seq Analysis

Data (40) was downloaded from GEO and loaded into a Seurat (v4.4.0) (41) object. The DotPlot function was used for Figure 3d and Figure S3e. To characterize fibroblast subpopulations, we identified highly variable genes, scaled expression data, and then computed PCA and UMAP visualizations on the fibroblast cells using Seurat. Gene signature scores from Schütz et al. were calculated with the AddModuleScore_UCell function from the UCell R package (42) (v2.4.0).

For CIBERSORTx deconvolution (run on web portal November 2023), cell subpopulation signatures were used to create a signature matrix representing each cell type. For immune deconvolution, the LM22 signature set provided by CIBERSORTx was used. The bulk expression matrix used as input for the SCC and KA mixtures consisted of VST-normalized expression data. Any negative values in the matrix were set to zero.

### GTEx Analysis

Gene-level count data and sample metadata were downloaded from the GTEx website. This count matrix was used as input for DESeq2 and differential expression tests between sun-exposed and sun-protected skin were performed with adjustment for sex and age. Log2 fold changes were computed using the apeglm (v1.16.0) (45) R package. The top 150 upregulated and downregulated genes with at least 100 mean base expression were included in the EvP signature. EvP scores were calculated using the same method as DvP scores.

### Statistical Methods

Kruskal-Wallis and Wilcoxon rank sum tests were used to compare values between groups. Spearman’s rho was used to calculate correlations unless otherwise specified. Statistics were computed using R (v4.3.1).

## Results

### Curation of publicly available SCC RNA-Seq datasets

A search for publicly available RNA-Seq samples of SCC, AK, IEC, KA, and/or NS identified 12 studies published in 2016–2023 with an average of 32 samples that met our initial inclusion criteria (**Table 1**) (21–32). Raw FASTQ data from these studies were downloaded and uniformly processed using STAR (46) for sequence alignment and RSEM (47) for read quantification; reads were aligned to the GENCODE v38 reference. A quality control filter was applied after alignment that required samples to have >60% mapped reads and >5 million total mapped reads. Samples that did not meet this criterion were removed from further analysis (**Figure S1a)**. Six additional samples demonstrated aberrant expression patterns that suggested possible misclassification and were also removed from our combined cohort. This left 306 samples from 11 studies with an average of 65 million reads per sample (**Figure S1b**).

**Table 1.** Publicly available SCC RNA-Seq studies considered in this analysis.

| **Author** | **Journal** | **Year** | **Samples** |
| --- | --- | --- | --- |
| Chitsazzadeh et al. | Nature Communications | 2016 | 26 |
| Hoang et al. | PeerJ | 2017 | 29 |
| Lukowski et al. | Journal Investigative Dermatology | 2018 | 25 |
| Cho et al. | Science Translational Medicine | 2018 | 18 |
| Lee et al. | Journal Investigative Dermatology | 2018 | 6 |
| Wan et al. | Genes & Diseases | 2019 | 70 |
| Das Mahapatra et al. | Scientific Reports | 2020 | 16 |
| Li et al. | EBioMedicine | 2020 | 16 |
| Vandiver et al. | Experimental Dermatology | 2021 | 23 |
| Srivastava et al. | Cancer Research | 2022 | 24 |
| Veenstra et al. | Journal Investigative Dermatology | 2023 | 20 |
| Bailey et al. | Nature Communications | 2023 | 110 |

Our final analysis cohort consisted of 73 NS, 46 AK, 16 IEC, 13 KA, and 147 SCC. Eleven samples labeled as AK/IEC or AK/IEC/SCC were also included (48) (**Table 2**). To the best of our knowledge, this cohort represents the largest collection of publicly available SCC RNA-Seq and includes the 70 SCC profiled by Bailey and colleagues, which is the next largest study (32). By comparison, a survey of SCC microarray studies showed an average of 15 SCC per study **(Table 3)**, about 10x smaller than our combined cohort. The large sample size of our study thus provides enhanced statistical power for unbiased differentially expressed gene (DEG) detection and the identification of consensus genes that are robust to inter-study variation.

**Table 2.** Number of each sample type after removing low quality samples. “Other” includes samples that were labeled as AK/IEC or AK/IEC/SCC.

| Sample Type | Sample Size |
| --- | --- |
| NS | 73 |
| AK | 46 |
| IEC | 16 |
| KA | 13 |
| SCC | 147 |
| Other | 11 |

**Table 3.** SCC microarray studies considered in this study.

| **Author** | **Journal** | **Year** | **Number of SCC** |
| --- | --- | --- | --- |
| Dooley et al. | Biochemical and Biophysical Research Communications | 2003 | 4 |
| Nindl et al. | Molecular Cancer | 2006 | 5 |
| Haider et al. | Journal Investigative Dermatology | 2006 | 8 |
| Kathpalia et al. | Journal of Dermatology | 2006 | 5 |
| Wenzel et al. | International Journal of Cancer | 2008 | 42 |
| Hudson et al. | Molecular Carcinogenesis | 2010 | 12 |
| Padilla et al. | Archives of Dermatology | 2010 | 15 |
| Sand et al. | Journal Dermatological Science | 2012 | 7 |
| Hameetman et al. | BMC Cancer | 2013 | 15 |
| Prasad et al. | Modern Pathology | 2013 | 6 |
| Brooks et al. | Journal of Clinical Investigation | 2014 | 10 |
| Lambert et al. | British Journal of Cancer | 2014 | 30 |
| Mitsui et al. | Journal Investigative Dermatology | 2014 | 5 |
| Sand et al. | Journal Dermatological Science | 2016 | 3 |
| Warren et al. | Scientific Reports | 2016 | 24 |
| Wei et al. | Journal of Cancer | 2018 | 40 |
| Burgo et al. | JCI Insight | 2018 | 13 |
| Qin et al. | Translational Cancer Research | 2021 | 32 |
| Hu et al. | BMC Genomics | 2022 | 6 |

Following sample-level quality control measures, the count data was normalized with DESeq2 (49). Principal component analysis (PCA) revealed study-specific clustering that indicated the presence of batch effects driven by study membership (**Figure 1a**), prompting batch correction using the removeBatchEffect function in the R limma package (50). Visual inspection of the post-correction PCA plots demonstrated that samples were no longer clustered by study but rather by sample type, arranging along the PC1 axis from normal skin to SCC (**Figure 1b**). To further validate this axis of neoplastic transition, we assessed the Differentiation-Progenitor-like (DvP) signature (32) in the batch-corrected expression profiles. A strong correlation between PC1 and DvP score was observed, with SCC demonstrating lower DvP scores than NS consistent with a more progenitor-like undifferentiated cell state (**Figure 1c**). By contrast, AK and IEC displayed intermediate DvP scores that were concordant with their respective positions along the SCC disease continuum. Visualization of samples with Uniform Manifold Approximation and Projection (UMAP) (51) after batch correction also showed that normal and malignant tissues were located in distinct clusters (**Figure S1c**). Taken together, this confirmed successful removal of study-level noise while retaining important biological variation.

**Figure 1.**
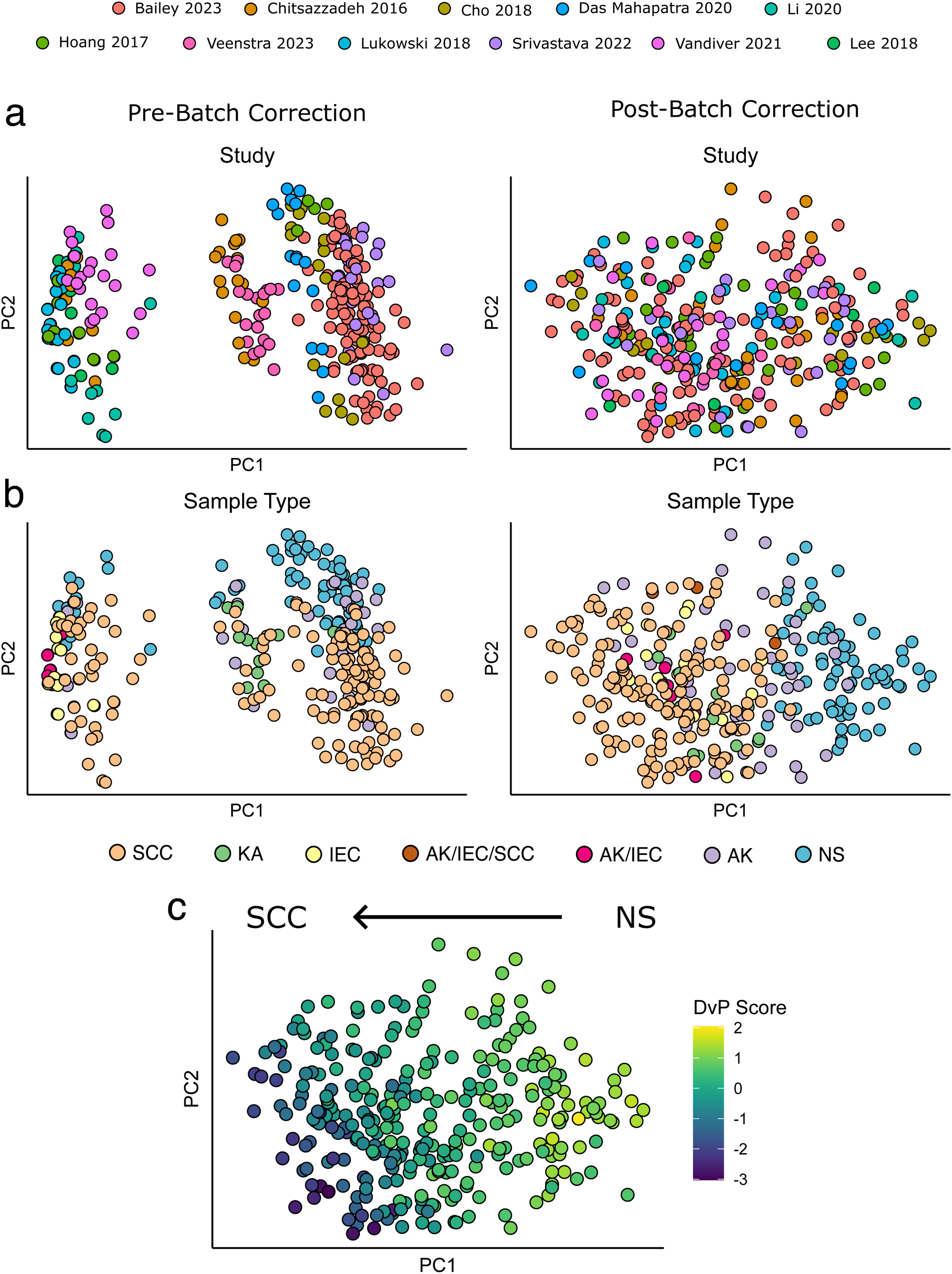
Analysis of publicly available SCC RNA-Seq studies. (a,b) Principal component plots showing study membership and sample type for each sample before and after batch correction with limma’s removeBatchEffect function. Samples display study-specific clustering prior to batch correction. PCA analysis was performed using the top 500 most variable genes. (c) PCA plot with DvP scores.

### Individual study DEGs reveal lack of consensus genes

Differential expression estimates are known to vary between studies (52,53) and a previous meta-analysis of SCC microarray datasets identified minimal DEG overlap between reports (35). To determine whether RNA-Seq studies of SCC display similar inconsistency in DEG detection, we performed differential expression analysis on each of the individual studies that collected both SCC and normal skin samples (n=9). A 5% false discovery rate (FDR) correction was used to provide a consistent analytical protocol to maximize comparability between studies. The number of DEGs varied widely (min 476 to max 17,021) (**Figure 2a**). Sample size was positively correlated with the number of DEGs detected per study but did not reach statistical significance (Pearson coefficient 0.47, p = 0.2) (**Figure S2a**). Thus, additional factors, such tumor heterogeneity or variable sample processing, may contribute to differences in DEG detection.

**Figure 2.**
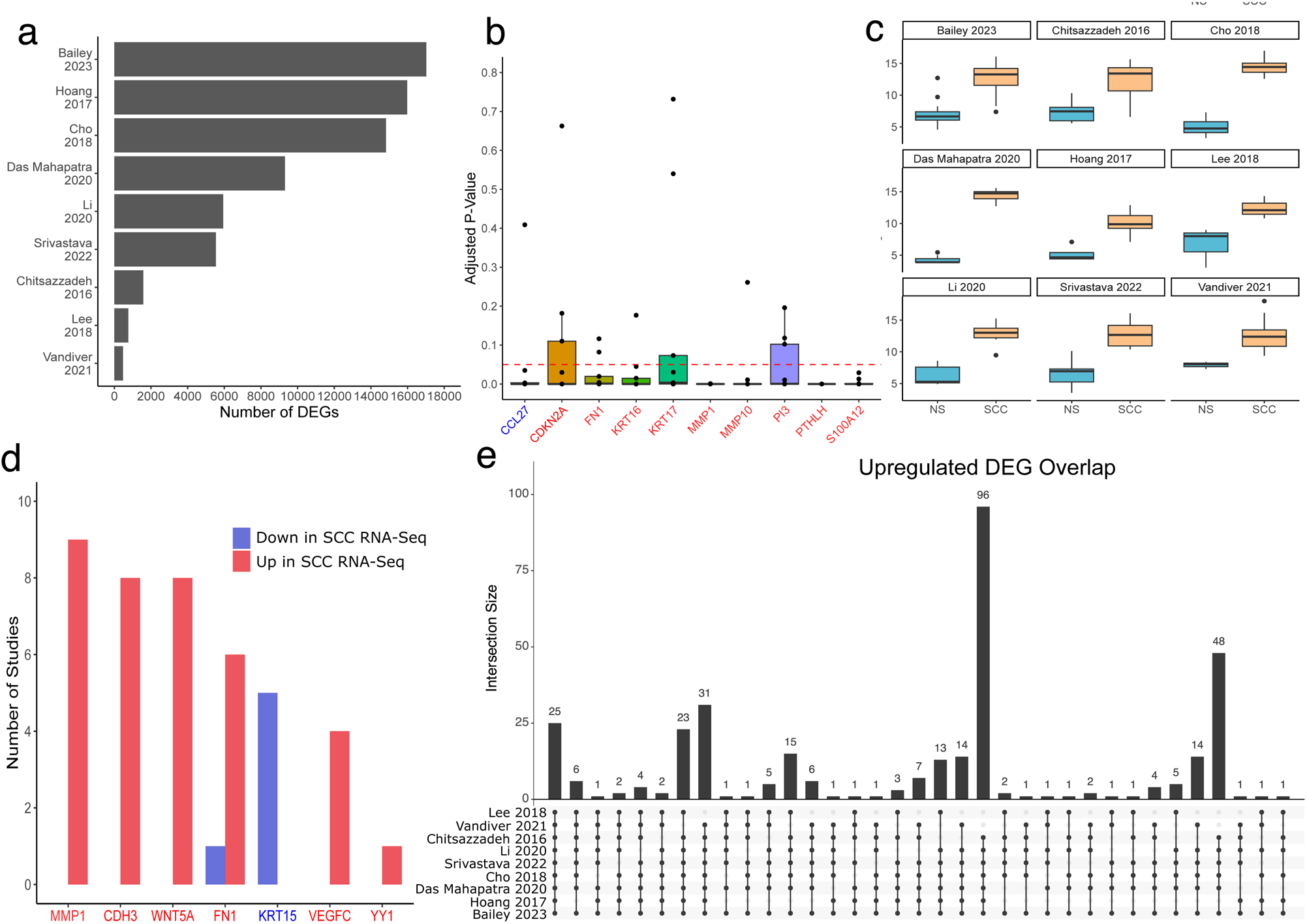
Comparing differentially expressed genes between individual studies reveals large inter-study heterogeneity. (a) Number of differentially expressed genes detected by DESeq2 when comparing normal skin vs SCC in each individual study. DEGs detected with 5% FDR using Benjamini-Hochberg correction. (b) Boxplots showing adjusted p-values of canonical SCC genes in each study. Each dot is the adjusted p-value for one study. Red dotted line indicates 5% FDR threshold. (c) Boxplots showing *MMP1* expression across studies with SCC and normal skin samples. (d) Barplot displaying the number of studies where genes identified by Van Haren et al. (35) were statistically significant. Gene names in red are predicted to increase in SCC and those in blue are predicted to decrease. (e) UpSet plot showing overlap of DEGs upregulated in SCC compared to normal skin between studies. The leftmost bar shows the number of upregulated DEGs detected in all studies.

We next surveyed previously published SCC microarray studies (2,3,5,7,10,11,13,17,35,54) and identified 10 genes with robust expression changes in two or more studies **(Supplementary Data 1)**. Examination of these canonical SCC genes in each of the nine RNA-Seq studies showed that only 3 of 10 genes (*PTHLH*, *MMP1*, and *S100A12*) were significantly changed in the expected direction (**Figure 2b and 2c**). An additional group of DEGs identified in a meta-analysis of non-melanoma skin cancer microarray studies (35) showed even less concordance, with only 1 of 7 genes (*MMP1*) detected by all RNA-Seq studies (**Figure 2d**). These findings highlight the variability of DEGs between studies and demonstrate that even well-known SCC genes do not always replicate in every report.

We then examined the degree of DEG overlap between studies in an unbiased manner by quantifying the number of shared upregulated and downregulated genes in SCC (**Figure 2e and S2b**). Only 25 upregulated genes and 8 downregulated genes were shared by all 9 studies; however, EnrichR (55) overrepresentation analysis performed on DEGs identified in at least 7 studies highlighted several pathways previously implicated in SCC, such as cell cycle genes and epithelial-mesenchymal transition (EMT) (56), (**Figure S2c**). Thus, pathway analysis tools can correctly identify disease-relevant pathways despite the low DEG overlap associated with smaller sample sizes.

### Pooling studies identifies consensus DEGs and pathway changes in SCC tumorigenesis

Utilizing our uniformly processed dataset, we conducted negative binomial regression analysis using DESeq2 and removed study-specific confounding by adjusting for study membership. This approach identified 14,385 consensus DEGs that were differentially expressed in NS vs SCC at a 0.1% FDR, with 6,862 DEGs having a Log2 fold change magnitude larger than 1 (**Figure S3a**). We also defined a set of genes with aberrant expression in the transition from NS to AK and the subsequent progression from AK to SCC (**Figure 3a-b**). In contrast to the individual study analysis findings described above, all canonical SCC genes changed significantly with the expected directionality in the pooled analysis **(Figure 3c**). *MMP10*, *PTHLH*, and *MMP1* showed an additional increase in expression during the transition from IEC to SCC. These genes are known to be expressed by tumor-specific keratinocytes (TSKs) (40) that localize to the invasive edge of SCC tumors, suggesting TSKs are increased in frank carcinoma compared to preinvasive IEC. Pathways previously implicated in SCC, such as interferon signaling, EMT, and lipid metabolism (6,32,56–58), were identified by enrichment analysis and overlapped with pathways found from individual-study analysis (**Figure 3d**).

**Figure 3.**
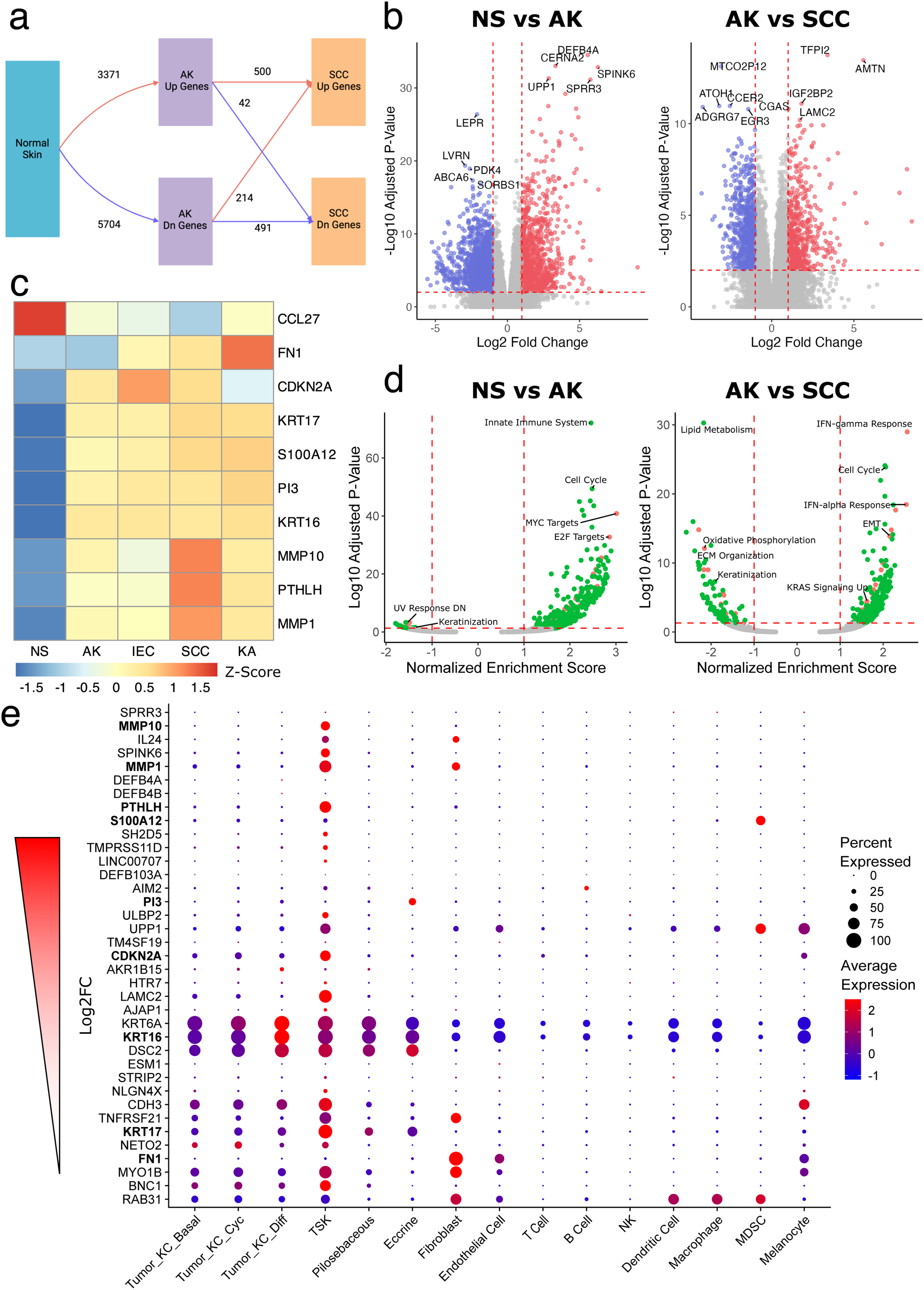
Pooling studies identifies consensus differentially expressed genes and pathway changes in the progression from normal skin to SCC. (a) Diagram with the number of DEGs in the transition from NS to AK and the subset of those genes that show additional changes in the progression from AK to SCC. 1% FDR threshold was used to identify DEGs. (b) Volcano plots showing genes differentially expressed in NS compared to AK and AK compared to SCC. Dotted red line indicates the 1% FDR threshold. (c) Expression of canonical SCC genes across transition from NS to SCC. Z-scores computed from VST-normalized expression data. All genes are predicted to be decreased in NS compared to SCC except for *CCL27*, which is predicted to be increased. (d) Volcano plots showing enrichment of Hallmark and Reactome gene sets in the same comparisons as (b). Dotted red line indicates 5% FDR threshold. (e) Expression of consensus genes upregulated in SCC in different cell populations defined by scRNA-seq from SCC tumors. Bolded genes are canonical SCC genes.

Recent work by our group suggests RET activation plays a role in SCC pathogenesis and demonstrates the therapeutic potential of RET inhibition in this malignancy (30). As further support of these findings, consensus DEGs included the RET ligand *ARTN* as well as RET biomarkers *DUSP6* and *SPRY4* (59), which were increased in the progression from NS to SCC in the combined cohort (**Figure S3b**). A signature of computationally predicted downstream RET transcriptional targets from head and neck squamous cell carcinoma (60) was similarly enriched in SCC **(Figure S3c)**, consistent with our prior studies in smaller cohorts and confirming RET activation in this cancer.

To gain cellular context for our consensus DEGs, we next plotted the expression of our top upregulated and downregulated candidates in different cell populations using single-cell RNA-Seq (scRNA-seq) data from normal skin and SCC (40) (**Figure 3e and S3d**). Most of the upregulated genes were expressed in TSKs; however, *FN1* and *S100A12*, which were also canonical SCC genes, were primarily expressed in fibroblasts and myeloid derived suppressor cells (MDSCs). This analysis provides cell-type context that informs whether DEGs are tumor-intrinsic or extrinsic and can be used when designing model system experiments to study these genes.

Analysis of DNA sequencing data can nominate potential oncogenes or tumor suppressor genes (TSGs). To determine the extent to which RNA-Seq expression data corroborates these predictions, we examined expression patterns of 30 previously reported mutational drivers of SCC (61) (**Figure S3e**). Among known TSGs, *TP53* and *NOTCH2* showed decreased expression in SCC suggesting activation of nonsense-mediated mRNA decay by cancer-associated mutations; however, other TSGs (*PTEN* and *NOTCH1*) demonstrated no significant change in expression. Moreover, expression level changes in the direction opposite of predicted were observed for both TSGs (*CDKN2A* and *FAT1*) and oncogenes (*CCND1*). Increased expression of *CDKN2A* in SCC has been previously noted (54) and reduced tumor purity from stromal cell contamination has been suggested (62); however, our findings indicate *CDKN2A* is predominantly expressed by TSKs (**Figure 3e**). These results demonstrate that expression changes do not necessarily mimic mutational consequences in driver genes and highlight the limitations of interpreting driver gene status from expression data alone.

The inclusion of 13 KA samples in our pooled cohort enabled comparison of these well-differentiated squamoproliferative lesions to SCC. E2F target expression was significantly decreased in KA compared to SCC, suggesting less cellular proliferation (**Figure S4**). The reduced proliferative capacity of KA was further supported by increased NOTCH signaling and lower levels of MYC activation (**Figure S4**). These findings agree with previous reports demonstrating inactivating *NOTCH* mutations in growing KA but not regressing lesions (63,64) and suggest that the KA in our cohort were collected during the terminal differentiation/involution phase rather than the initial growth phase.

### Bulk deconvolution provides insight into the cellular composition of SCC subtypes

To further leverage previously published scRNA-seq data, we next applied CIBERSORTx (65) to infer the cellular composition of different SCC subtypes. We focused on SCC and KA as these lesions are expected to have the highest proportion of malignant cells that are well-represented by available scRNA-seq data. Deconvolution revealed KA contain more differentiated tumor cells and fewer TSKs compared to SCC; these findings are in line with the shift towards terminal differentiation observed during KA regression (66) and their benign clinical behavior (**Figure 4a**). By contrast, SCC arising in the setting of recessive dystrophic epidermolysis bullosa (RDEB) in our cohort (n=8) showed equivalent levels of differentiated tumor cells and higher levels of TSKs than sporadic tumors, consistent with their clinically aggressive phenotype (67) (**Figure 4b**). For both of these comparisons, the basal and cycling tumor cell populations were unchanged between SCC subtypes (**Table 4-5**).

**Figure 4.**
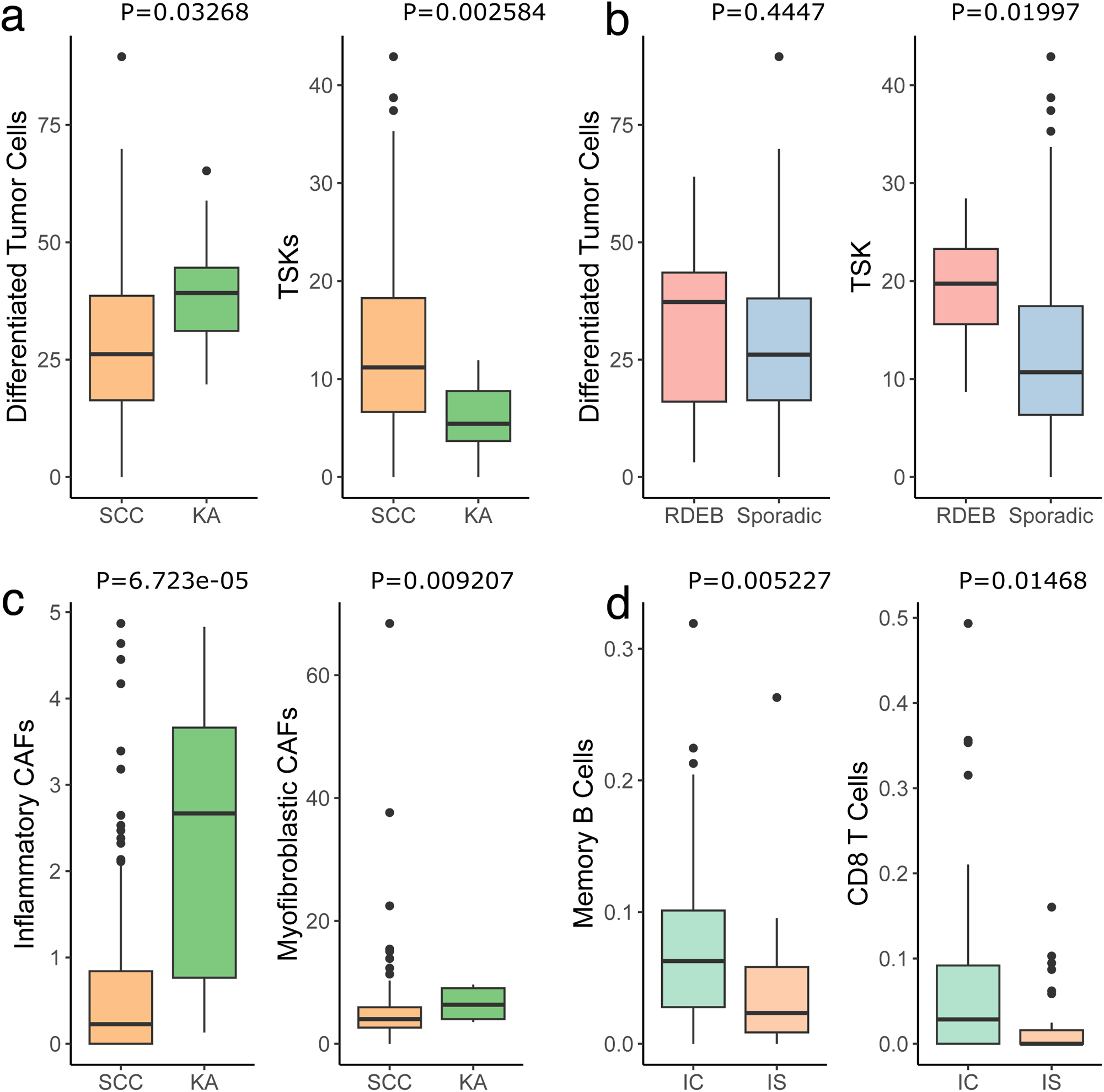
Bulk deconvolution provides insight into the cellular composition of SCC subtypes. (a,b) Boxplots showing levels of differentiated tumor cells and TSKs in SCC vs KA and RDEB-SCC vs. sporadic SCC. (c) Estimated levels of inflammatory and myofibroblastic CAFs in SCC vs KA samples. (d) Estimated levels of memory B cells and CD8 T cells in immunocompetent (IC) vs immunosuppressed (IS) SCC. P-values calculated using the two-sample Wilcoxon rank sum test.

**Table 4.** Tumor stromal subpopulations in SCC vs KA. P-values calculated with Wilcoxon rank sum test.

| Cell Type | Median SCC | Median KA | P-value |
| --- | --- | --- | --- |
| Basal Tumor Cells | 0 | 0 | 0.48787414 |
| Cycling Tumor Cells | 0 | 0 | 0.12217932 |
| Differentiated Tumor Cells | 26.1607628 | 39.2035544 | 0.03267985 |
| TSK | 11.1839328 | 5.43454125 | 0.00258391 |
| Myofibroblastic CAFs | 4.00465454 | 6.35801996 | 0.00920744 |
| Inflammatory CAFs | 0.22599365 | 2.66922899 | 6.72E-05 |

**Table 5.**
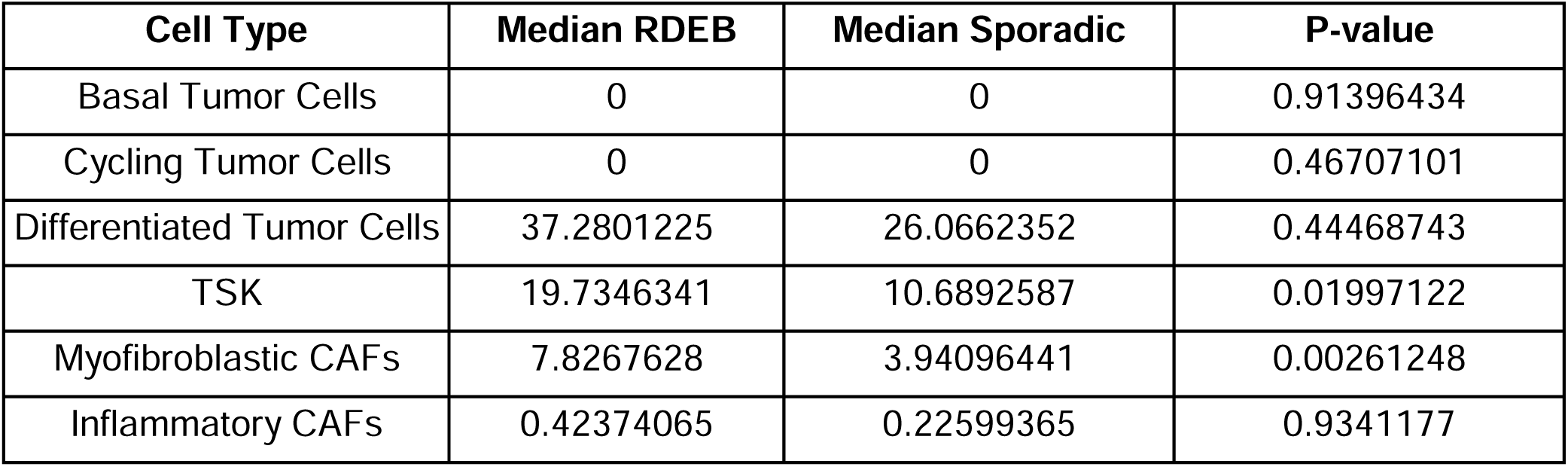
Tumor stromal subpopulations in RDEB-SCC vs sporadic SCC. P-values calculated with Wilcoxon rank sum test.

| Cell Type | Median RDEB | Median Sporadic | P-value |
| --- | --- | --- | --- |
| Basal Tumor Cells | 0 | 0 | 0.91396434 |
| Cycling Tumor Cells | 0 | 0 | 0.46707101 |
| Differentiated Tumor Cells | 37.2801225 | 26.0662352 | 0.44468743 |
| TSK | 19.7346341 | 10.6892587 | 0.01997122 |
| Myofibroblastic CAFs | 7.8267628 | 3.94096441 | 0.00261248 |
| Inflammatory CAFs | 0.42374065 | 0.22599365 | 0.9341177 |

We then extended these findings further by defining populations of inflammatory (iCAF) and myofibroblast-like (myCAF) cancer-associated fibroblasts (CAF) in SCC subtypes using recently published signatures (68) (**Figure S5a and S5b**). Exploration of these fibroblast subpopulations in different SCC subtypes revealed higher levels of iCAFs in KA compared to SCC, which may reflect the prominent mixed inflammatory infiltrate often present in these lesions (**Figure 4c**). myCAF expression was also increased in KA, and we note that stromal expression of one of the top myCAF marker genes, *WNT5A*, is known to induce differentiation and regression in skin tumors (69). Furthermore, Wnt5a can inhibit canonical Wnt signaling, which may promote KA regression (66). Interestingly, myCAFs, but not iCAFs, were also increased in RDEB-SCC compared to spontaneous SCC (**Figure S5c and Table 5**). The distinct clinical differences between KA and RDEB-SCC suggest that iCAFs may modulate the net effect of myCAFs to encourage either tumor regression or promotion.

Given the increased incidence of SCC in immunosuppressed (IS) individuals compared to immunocompetent (IC) persons (70,71), we next sought to explore the cellular composition of SCC in these patients to search for differences that might explain their different cancer susceptibilities. Deconvolution analysis of 77 IC and 30 IS SCCs (**Table 6**) identified a significant reduction in memory B cells in the latter, confirming recent findings (72). Our effort also revealed fewer CD8 T cells in SCC arising in IS patients, which is consistent with the known mechanisms of action of many immunosuppressive medications (**Figure 4d**). T follicular helper cells, activated dendritic cells, and activated mast cells showed marginal differences between immune states. Other immune cell types, including CD4 T cells, macrophages, and NK cells, were comparable between IC and IS SCC (**Table 7**).

**Table 6.** Details on immunosuppression for tumors in IC vs IS CIBERSORTx analysis.

| Type | Number of Patients |
| --- | --- |
| Graft Versus Host Disease | 1 |
| Cardiac Transplant | 2 |
| Immunosuppressed, no specifics | 5 |
| Renal Transplant | 22 |
| Immunocompetent | 77 |

**Table 7.**
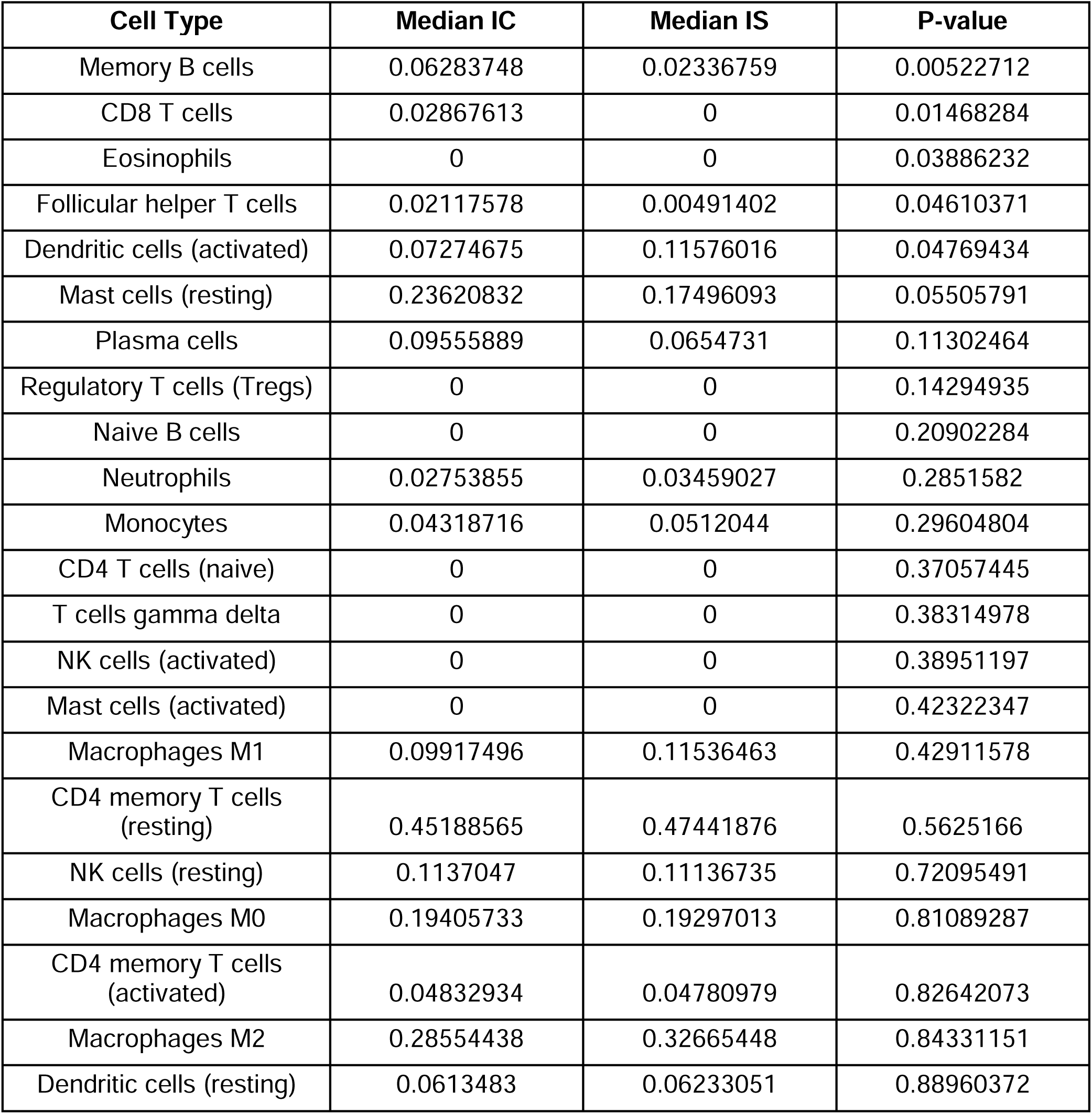
Immune cell subtype comparisons in IC vs IS patients. P-values calculated with Wilcoxon rank sum test.

### Unified fusion calling highlights lack of recurrent driver fusions

SCC is believed to be driven predominantly by UV-induced mutations (61,73), raising questions regarding the extent to which gene fusions contribute to SCC pathogenesis. Two fusions, EGFR-PPARGC1A and ADCK4-NUMBL, have been identified in SCC cell lines and confirmed in human samples via a targeted approach (33,34); however, to date an unbiased search for gene fusions in patient SCC samples has not been performed. Using the STAR-Fusion pipeline (74), we first sought to confirm the presence of EGFR-PPARGC1A and ADCK4-NUMBL fusions in our full cohort as well as identify any additional fusions. A total of 395 unique fusions were detected in patient-derived tissues and 52 unique fusions were called in SCC cell lines A431 and SCCIC1 (75,76). Fusions were detected in the majority of samples, with variation observed between studies and across sample types **(Figure S5ac)**. A431 cells had significantly more fusions than SCCIC1 cells or SCC tumors (22.7 vs 6 or 5.8 respectively, F-test P<2.2e-16), suggesting the A431 genome is more altered than most patient tumors. The EGFR-PPARGC1A fusion was successfully identified in all three replicates of A431 cells, consistent with prior reports and validating our use of STAR-Fusion. EGFR-PPARGC1A was not found in SCCIC1 cells and ADCK4-NUMBL was not detected in either cell line. Unexpectedly, neither fusion was identified in any of the 302 patient samples tested with STAR-Fusion, suggesting their prevalence may be lower than previously suggested or reduced tumor purity in bulk samples may confound fusion detection.

Surprised by the lack of previously studied fusions in patient samples, we turned our attention to novel SCC fusions. Of the 395 fusions found in patient samples, 22% (86 of 395) were detected in NS, suggesting they are less likely to be pathogenic. The remaining 78% (309 of 395) were present only in lesional tissue (“lesional fusions’’); however, most of these were not shared across sample types **(Figure S5d)** and only 6 fusions were shared between SCC cell lines and patient tumors **(Figure S5e)**. We next reasoned that fusion transcripts present in normal tissues from other organs (77,78) are less likely to be pathogenic, while fusions detected in tumors from The Cancer Genome Atlas (TCGA) (79,80) or that involve a COSMIC gene (81) are more likely to play a functional role. Using these criteria, 11% (33 of 309) of lesional fusions were present in other normal tissues. About 14% (42 of 309) of lesional fusions were present in TCGA tumors and 7% (21 of 309) had a COSMIC gene partner **(Figure S5f**), leaving the pathogenicity of most lesional fusions unclear.

We subsequently sought to prioritize the 276 lesional fusions not present in normal tissues by their recurrence (**Supplementary Data 2**). None of these candidates were present in more than 10% of SCC, with the top fusion, KRT6C-KRT127P, present in 9% (13 of 146). While KRT6C is significantly upregulated in SCC (82), the KRT6C-KRT127P fusion is not predicted to have a coding sequence, making its function less clear. The next hit, KRT5-KRT84, is predicted to produce a coding protein and was observed in 88% (14 of 16) of IECs, but only 8% (12 of 146) of SCC, suggesting a lack of positive selection. Among lesional fusions also present in TCGA tumors, the most recurrent hit, KRT16-KRT6A, is an interchromosomal in-frame fusion that occurs in 5% of SCC (8 of 146). An in-frame DMKN-KRT1 fusion was detected in 7% of KA (1 of 13) and is notable since DMKN is a marker of differentiation (83) and KA are highly differentiated neoplasms. Taken together, these findings suggest gene fusions are less likely to play a major role in SCC development.

### Chronic sun exposure signature from GTEx highlights UV-regulated genes that are involved in SCC pathogenesis

An open question in SCC biology is how UV radiation facilitates tumor development. To identify new genes regulated by chronic sun exposure that play a role in SCC carcinogenesis, we compared chronically sun-exposed skin (n=701) to sun-protected skin (n=604) in the GTEx skin dataset (77) and distilled a 300-gene sun-exposed vs sun-protected (EvP) signature to quantify the level of chronic sun exposure in human samples (**Figure 5a**). When used to score our pooled cohort, the EvP signature demonstrated higher levels of sun exposure in progressively dysplastic tissues (**Figure 5b**). EvP scores were most variable in NS, reflecting the inclusion of both sun-exposed and sun-protected samples in this group. We then used the EvP signature to compare RDEB vs sporadic SCC and observed lower scores in the former, reflecting the relatively minor role played by UV in RDEB-SCC (24,67) (**Figure 5c**).

**Figure 5.**
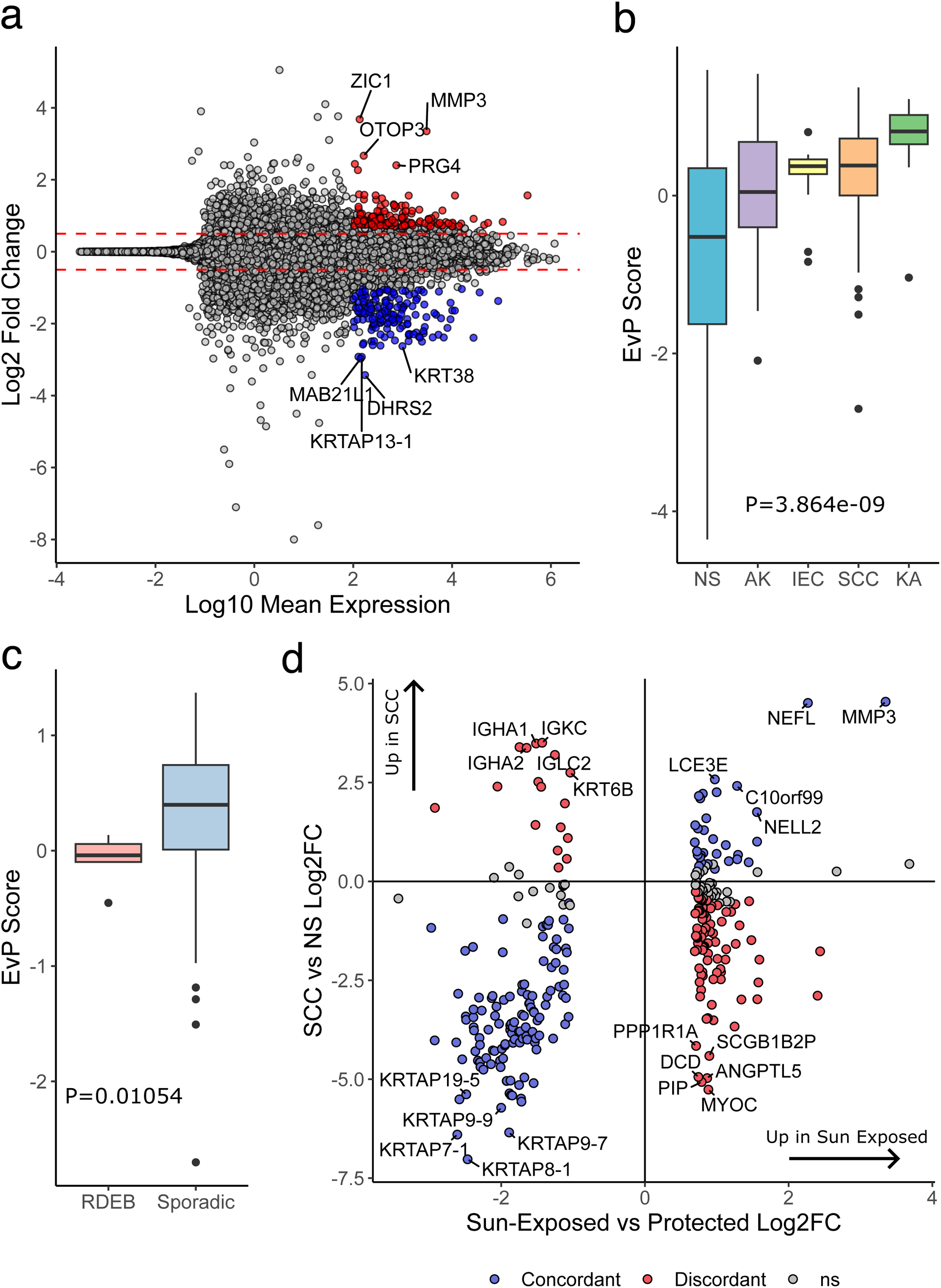
Chronic sun exposure signature from GTEx highlights UV-regulated genes that are involved in SCC pathogenesis. (a) MA plot showing DESeq2 log2 fold changes and average gene expression in GTEx sun-exposed and sun-protected skin samples. Highlighted genes are top hits from EvP signature. (b) EvP scores across the continuum of NS to SCC. Kruskal Wallis test used to compute P-value. (c) EvP scores in RDEB-SCC vs sporadic SCC. Wilcoxon rank sum test used to compute P-value. (d) Gene scores for EvP signature genes in normal skin vs SCC and sun-exposed vs sun-protected comparisons.

Next, we visualized the expression patterns of EvP genes in response to sun exposure and during the transition from NS to SCC (**Figure 5d**). Keratin-associated proteins (KRTAPs) were downregulated in both sun-exposed skin and SCC, suggesting UV radiation may impact hair follicles and consistent with their absence in SCC. Genes upregulated as a result of both sun exposure and malignant transformation included *MMP3* and *C10orf99*, with the latter recently shown to induce keratinocyte proliferation and inflammation (84). By contrast, immunoglobulin genes expressed by plasma cells were downregulated by sun exposure but upregulated in tumors, suggesting sun damage may attenuate plasma cell-mediated immunity that is re-activated when malignant keratinocytes reach their full neoplastic potential. These findings together nominate gene candidates regulated by UV that may functionally impact SCC development.

## Discussion

Individual RNA-Seq and microarray-based studies of cancers including SCC are often burdened by study-specific batch effects and smaller sample sizes that can limit the broad applicability of their findings (35,85). Our meta-analysis provides the largest cohort of publicly available SCC RNA-Seq samples to date and includes tissues at different stages along the progression of normal skin to invasive carcinoma, providing unmatched statistical power as well as new insight into SCC development.

The low number of DEGs shared across the cohorts pooled for this analysis highlights the need for a consensus list of DEGs in SCC, particularly one that includes information on genes involved in preinvasive stages. Our results confirm changes in genes and pathways previously reported in SCC and also identify previously unrecognized DEGs, such as elevated expression of the uridine phosphorylase encoded by *UPP1*, both in the progression of NS to AK as well as in the transition from AK to SCC. *UPP1*’s upregulation suggests increased uridine utilization may be important early in SCC development, consistent with its critical role in pancreatic cancer under glucose-restricted conditions (86). We also note that the stemness maintenance gene *TPFI2* is increased in SCC compared to AK, hinting that the transition from pre-neoplastic to malignant status may be marked by increased cancer stemness (87).

CIBERSORTx deconvolution was used to compare the cellular compositions of SCC, KA, and RDEB-SCC in our cohort. Consistent with our clinical understanding of these SCC subtypes, this analysis revealed RDEB-SCC have the most TSKs, providing further evidence that TSKs are responsible for more aggressive tumor phenotypes and worse outcomes. Deconvolution analysis also demonstrated lower levels of memory B cells in SCC arising from immunosuppressed patients compared to their immunocompetent counterparts, independently validating a recent finding by Thai and colleagues (72). More than half of Thai’s IS cohort was immunosuppressed due to hematologic malignancies, while our IS patients were predominantly renal transplant recipients, suggesting impaired memory B cell immunity in SCC is robust to different types of immunosuppression. This demographic difference may also explain why we identified lower levels of CD8 T cells – a key target in post-transplant immunosuppression – in SCC from IS individuals. Future studies augmenting B cell immunity, either through vaccinations or passive antibody transfers, should be conducted to evaluate whether this can improve SCC outcomes in IS patients.

Our unbiased and comprehensive search for fusion transcripts in primary human SCC and intermediate disease stages did not detect the two fusions previously reported in this malignancy. This difference is unlikely to be an artifact of our fusion detection algorithm as we were able to detect the EGFR-PPARGC1A transcript in A431 cells, where it was originally reported (33); however, stromal cells in primary specimens may mask fusion detection. Novel fusions were identified by STAR-Fusion at various stages along the continuum of SCC development, but these candidates were largely absent in TCGA tumors, did not involve COSMIC cancer genes, and demonstrated low rates of recurrence. Thus, our findings suggest gene fusions are not a common driver event in SCC.

We recognize that our analysis has important limitations. First, while we would have liked to correlate molecular features with patient outcomes, this information was not provided in most of the individual studies. Second, while we identified and removed study-specific batch effects and verified that batch correction retained important biological differences, we cannot be certain that all confounders were eliminated. Third, due to a paucity of large-scale SCC studies with paired mutation and expression data, our analysis only contained the latter, preventing comparison of genomic vs transcriptomic differences in SCC and precursor lesions. Future studies should strive to gather both types of data to provide better context to molecular findings.

In summary, by reprocessing and analyzing raw data from 11 studies, we have created a robust consensus list of DEGs that can be used by the research community, defined the cellular composition of different SCC subtypes, performed an unbiased search for gene fusions in SCC as well as related preinvasive tissues, and distilled a novel gene signature for quantifying chronic sun exposure. Our results provide an important resource for future studies directed at identifying new therapeutic targets in SCC.

## Acknowledgements

We thank Ankit Srivastava, Vanessa Lopez-Pajares, and Kenneth Tsai for helpful discussions. The results published here are in part based upon data generated by the TCGA Research Network: https://www.cancer.gov/tcga. The Genotype-Tissue Expression (GTEx) Project was supported by the Common Fund (https://commonfund.nih.gov/GTEx) of the Office of the Director of the National Institutes of Health, and by NCI, NHGRI, NHLBI, NIDA, NIMH, and NINDS.

## Data availability

Code to replicate the analyses in the paper is available at https://github.com/tjbencomo/scc-meta-analysis. Additional data supporting the findings of this study, including consensus differentially expressed genes and batch-corrected expression profiles, can be found on Zenodo at https://zenodo.org/records/10272679.

## Ethics statement

Ethics approval was not required for this study as data was collected from published studies in which informed consent was already obtained by the study investigators.

## Author Contributions

TB: Conceptualization (lead), data curation (lead), formal analysis (equal), investigation (equal), methodology (lead), visualization (equal), writing - original draft (equal), writing - review & editing (equal). CSL: Conceptualization (supporting), data curation (supporting), formal analysis (equal), funding acquisition (lead), investigation (equal), methodology (supporting), visualization (equal), writing - original draft (equal), writing - review & editing (equal).

---

**Figure S1.**
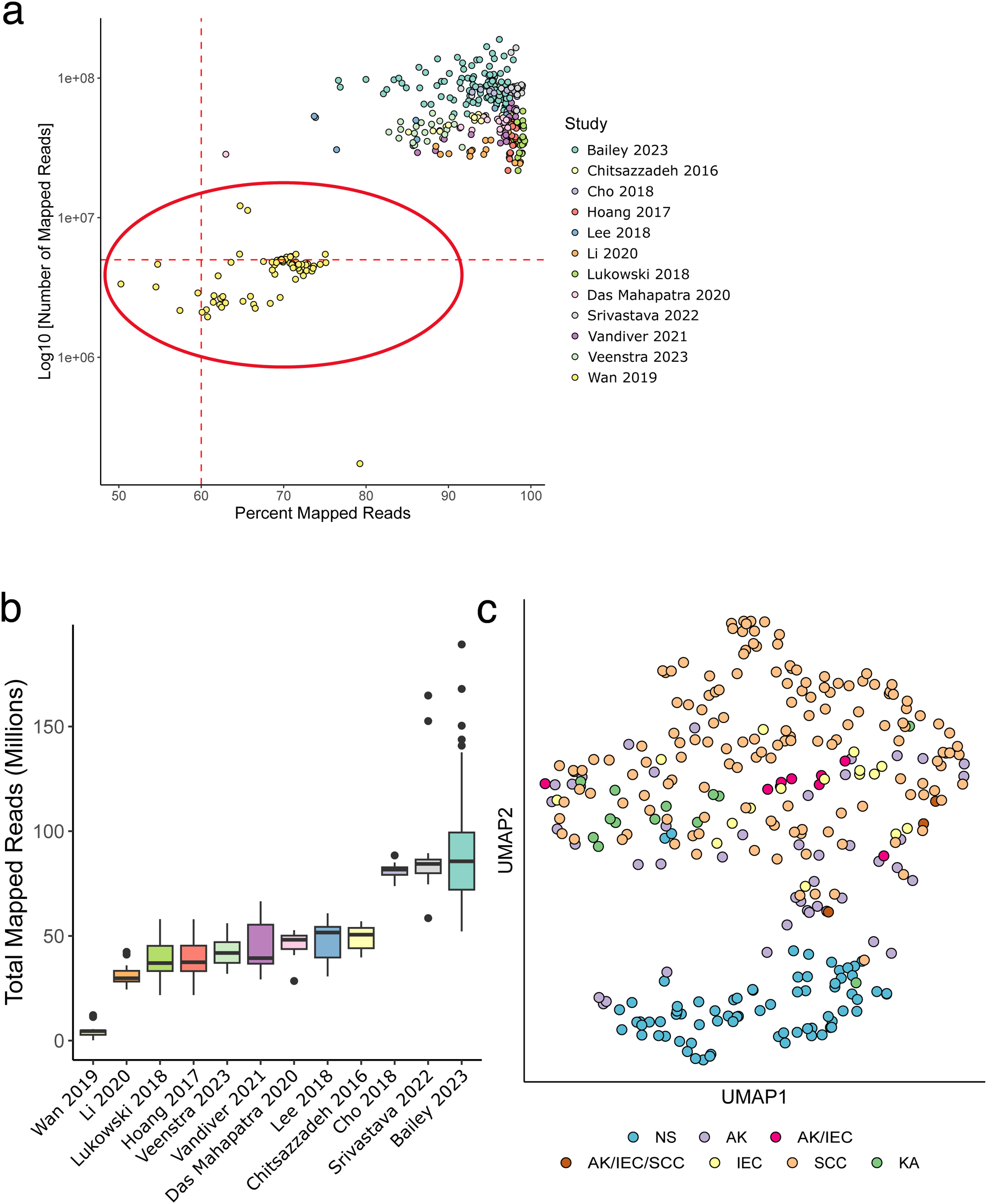
Sample quality control analysis. (a) Sequencing quality plot for all samples initially considered in the study. Red dotted lines are at 60% mapped reads and 5 million total mapped reads. Circled samples were removed from further analysis. (b) Mean number of total mapped reads per sample for each study. (c) UMAP showing samples cluster predominantly by lesion type.

**Figure S2.**
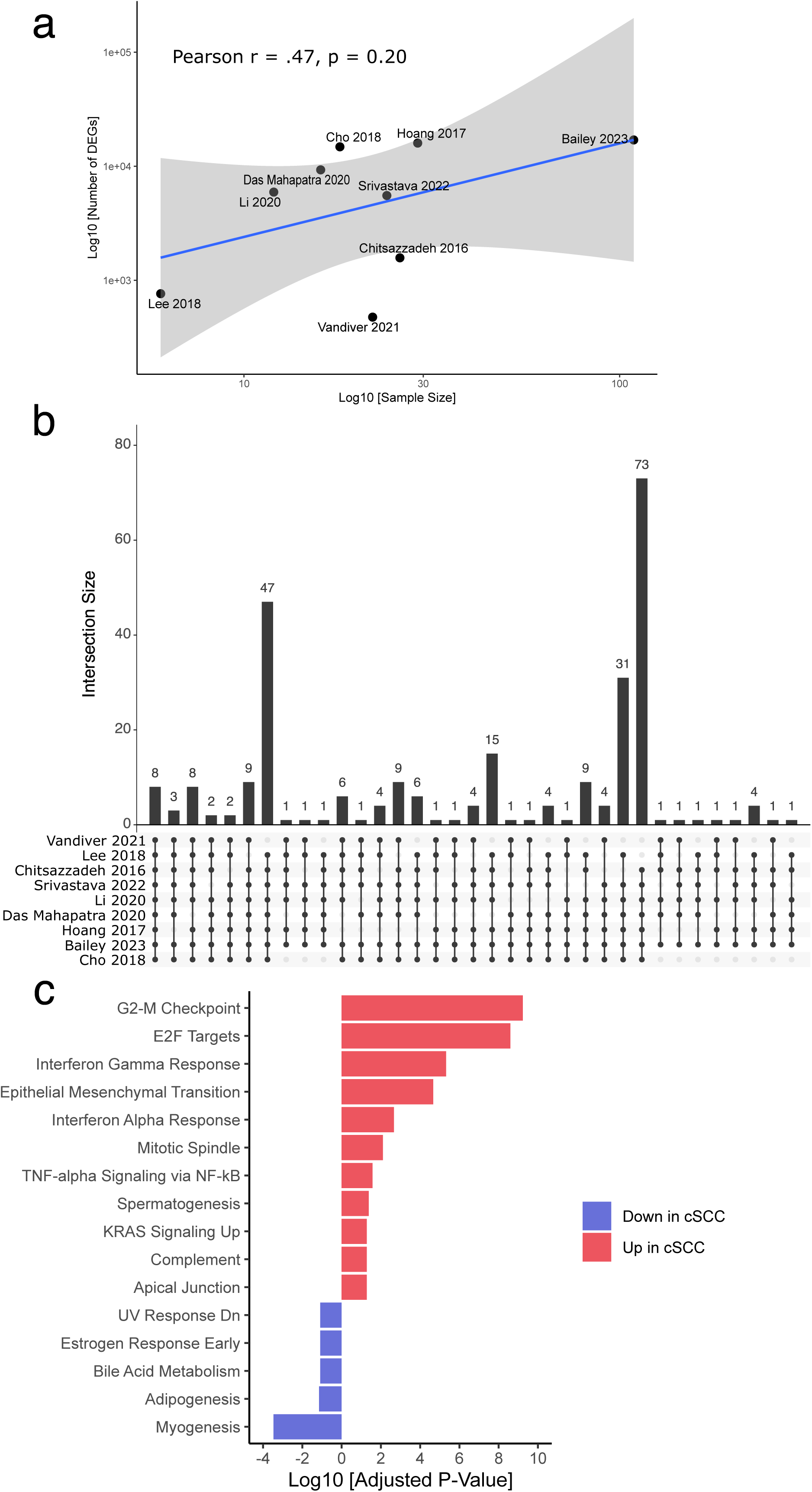
Analysis of DEG consistency. (a) Scatter plot comparing study sample size to number of DEGs detected in normal skin vs SCC. (b) UpSet plot showing overlap of DEGs downregulated in SCC compared to normal skin between studies. Leftmost bar shows the number of downregulated DEGs detected in all studies. (c) EnrichR results for upregulated and downregulated genes that were identified in 7 or more studies with a 10% FDR. Gene sets are from the 2020 MSigDB Hallmark collection.

**Figure S3.**
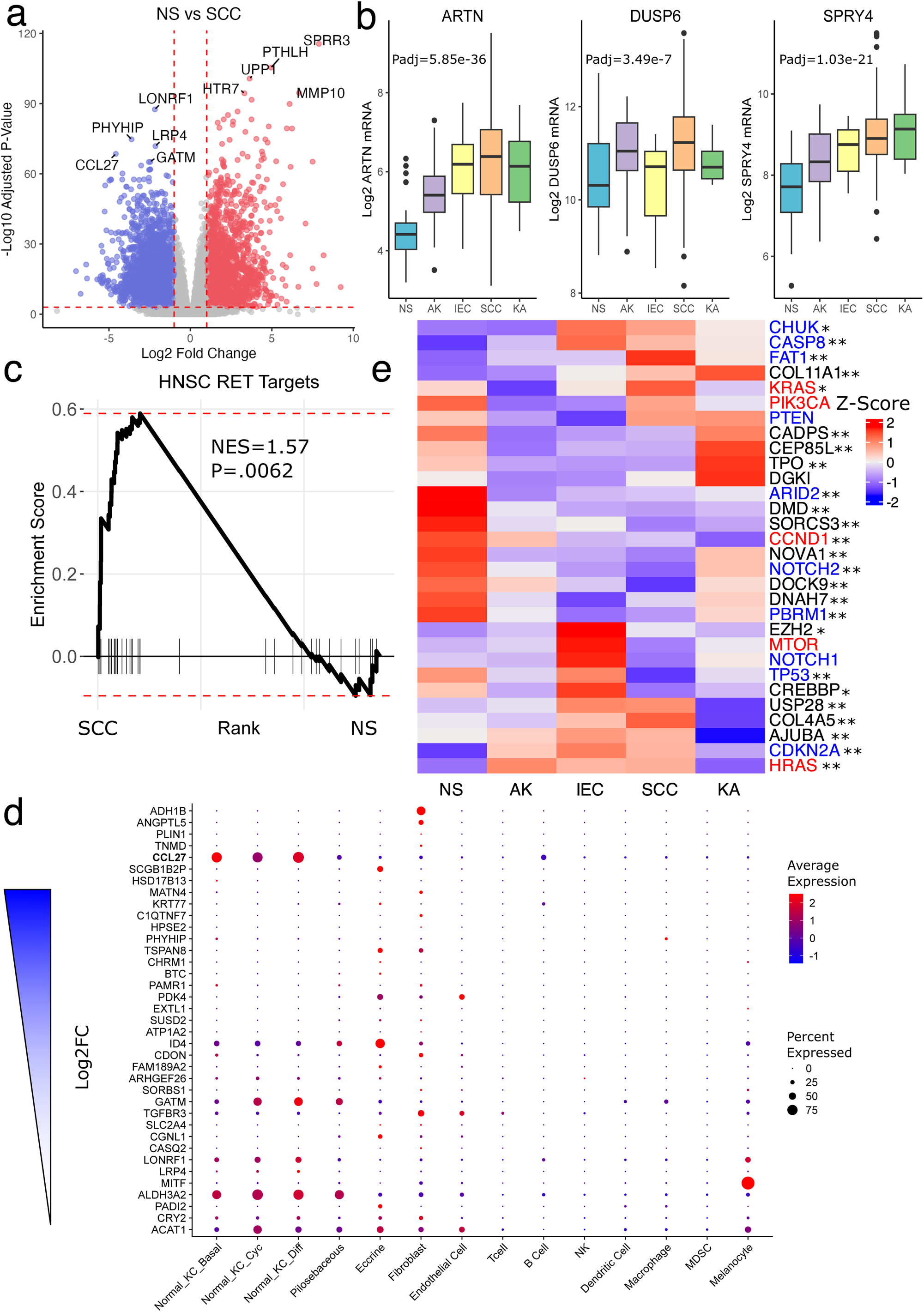
Pooled cohort DEG analysis. (a) Volcano plot showing DEGs from pooled normal skin vs SCC comparison. 0.1% FDR threshold used to identify DEGs. (b) Selected RET pathway genes that are upregulated in the transition from normal skin to SCC. P-values computed by DESeq2 likelihood ratio test. (c) GSEA enrichment of RET transcriptional targets in head and neck squamous cell carcinoma (HNSC) identified by ARACNe. (d) Expression of consensus genes downregulated in SCC in different cell populations defined by scRNA-seq from normal skin. Bolded genes are canonical SCC genes. (e) Heatmap displaying mRNA expression of genes determined to be under positive selection in SCC by at least one of three algorithms. Red genes are known oncogenes, blue genes are known tumor suppressors, and black genes have not been definitively classified. Asterisks represent statistical significance for NS vs SCC comparison via Wald test.

**Figure S4.**
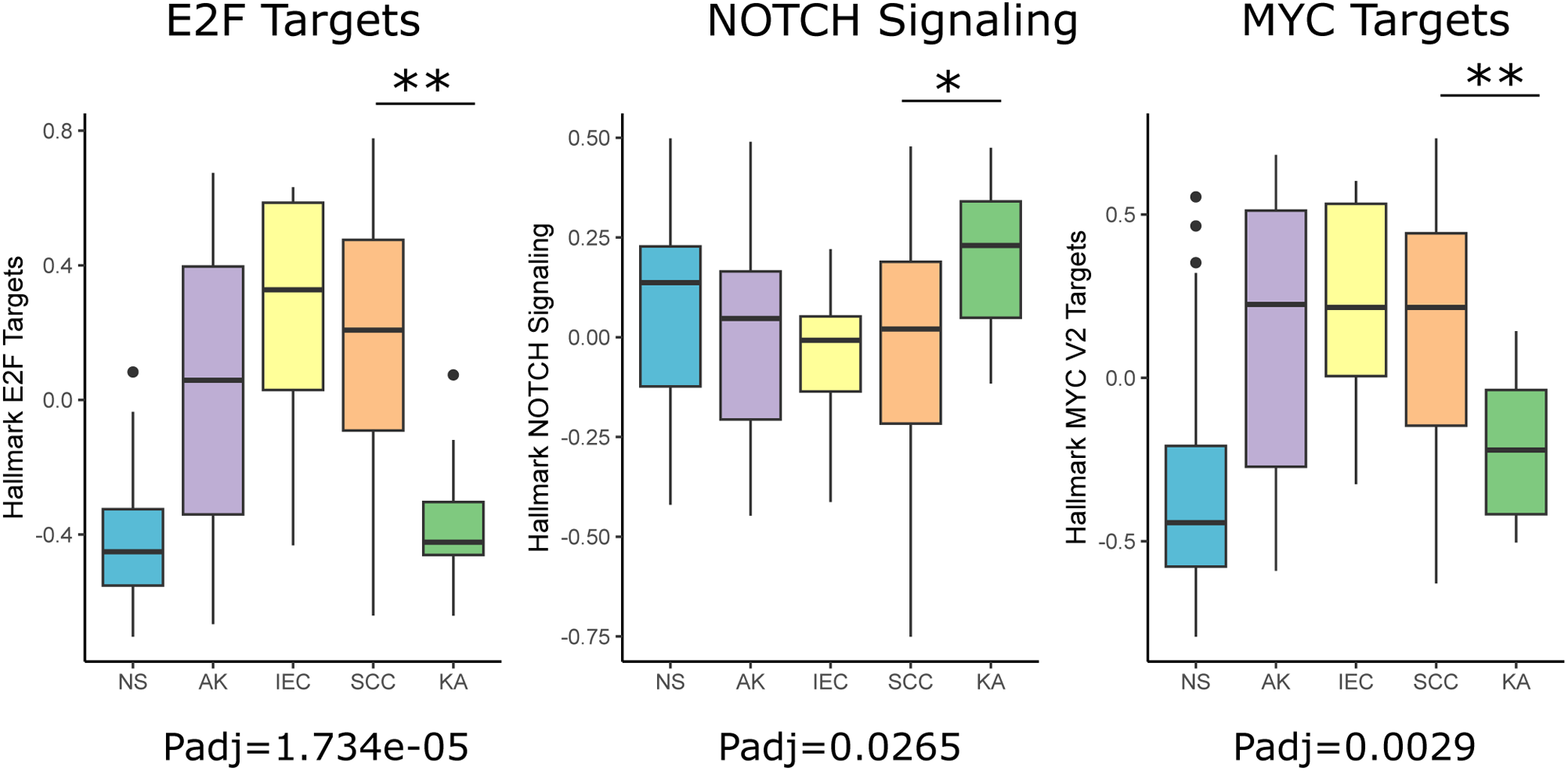
Pathway activity in keratoacanthomas. (a) Boxplots showing signature scores for E2F targets, NOTCH signaling, and MYC targets from MSigDB Hallmark collection. P-values from linear regression model with SCC and KA terms. Multiple comparisons correction used Benjamini-Hochberg method. ** P-value < .001, * P-value < .05.

**Figure S5.**
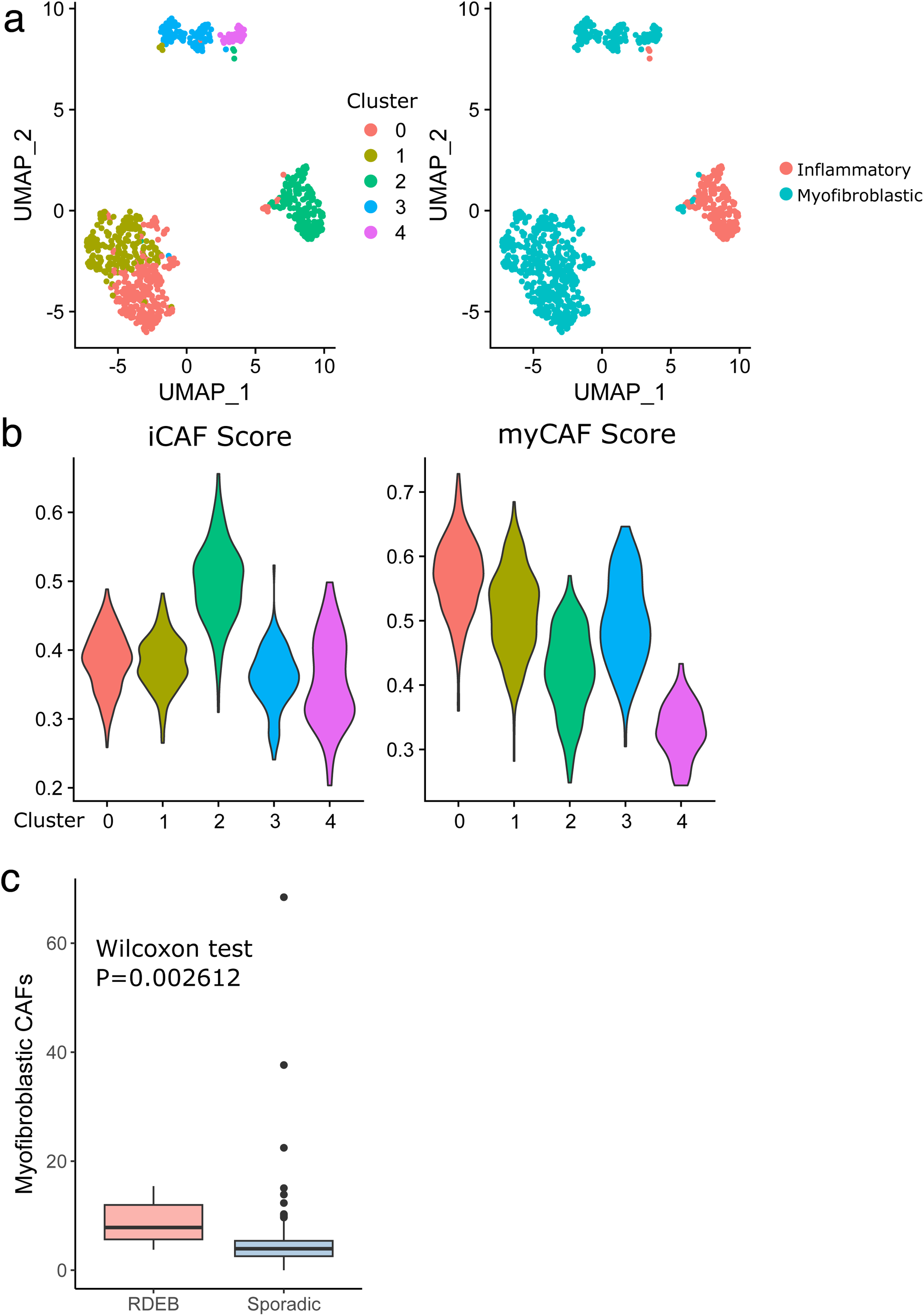
Lesional tissue deconvolution with CIBERSORTx. (a) UMAP plots showing fibroblast cells from Ji et al. (40). Cells are clustered using Seurat FindClusters (resolution=0.3). (b) iCAF and myCAF scores in Ji’s fibroblasts using signatures from Schutz et al. (68). Scores calculated using UCell. (c) Boxplot showing levels of myofibroblastic CAFs in RDEB-SCC vs sporadic SCC.

**Figure S6.**
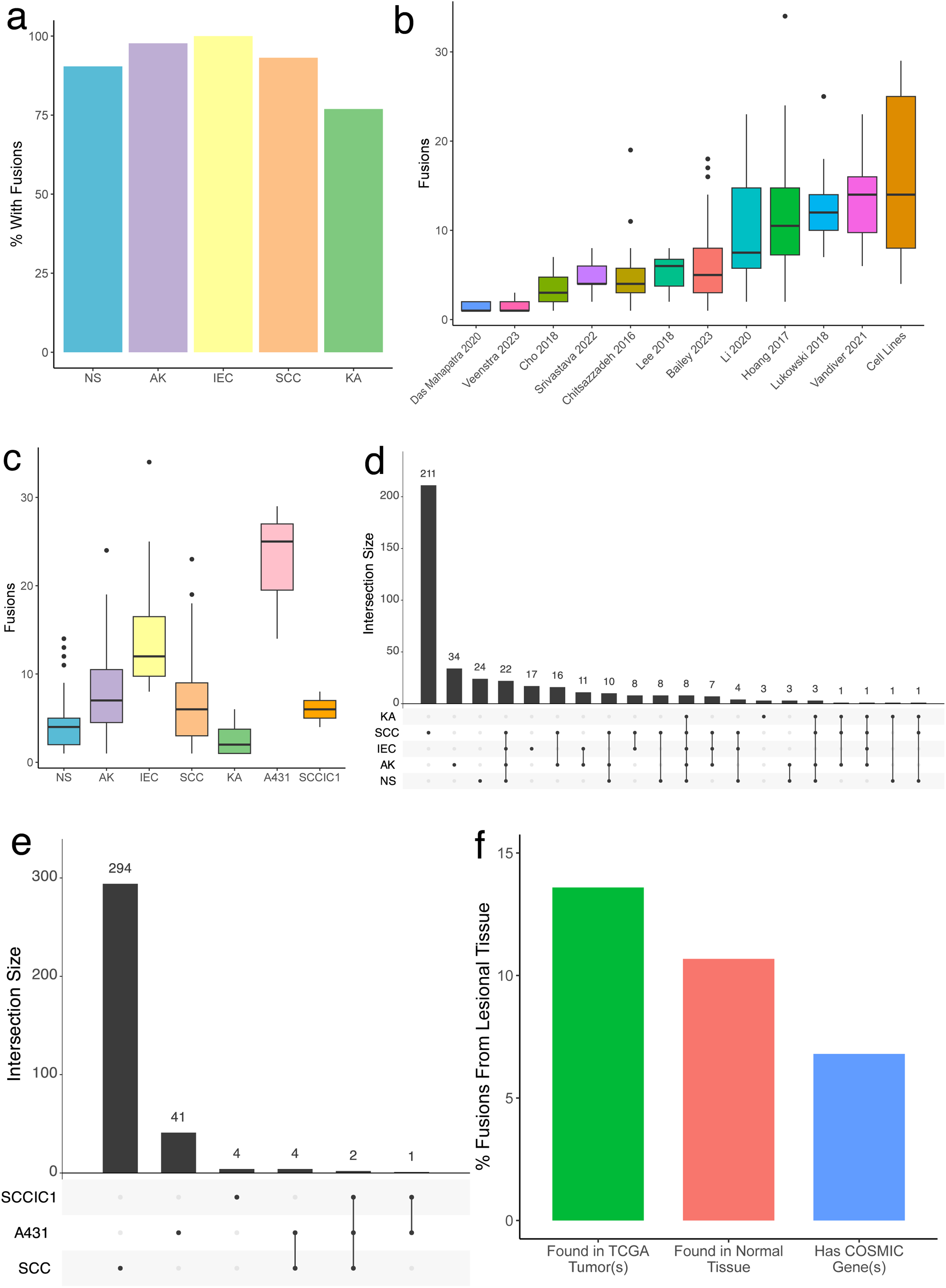
Characterization of gene fusions in NS and lesional tissue. (a) Percent of samples in each sample type with at least one fusion detected by STAR-Fusion. (b, c) Distribution of fusions per study cohort and sample type. (d) UpSet plot showing number of fusions shared between sample types. (e) Number of fusions shared between SCC samples and SCC cell lines. (f) Features of fusions found in non-normal skin. Fusions found in normal tissues were present in either GTEx or by Babiceanu et al. (78).

